# Geographical Distributions and Spatial Equilibrium in Historical Epidemics of the United States

**DOI:** 10.1101/486506

**Authors:** Stephen Coleman

**Affiliations:** Research Director and Professor, Retired Center for Applied Research and Policy Analysis Metropolitan State University St. Paul, Minnesota USA

**Keywords:** epidemics, geographical distribution, mathematical model, equilibrium, United States

## Abstract

This research examines the geographical distributions of several historical epidemics in the United States and investigates whether they reached a geographical equilibrium, however briefly. An equilibrium distribution over a geographical area, as the end state of a diffusion or spatial contagion process, has definitive mathematical properties. These permit qualitative and quantitative tests that may confirm an equilibrium and identify its characteristics. The analysis uses United States state-level data for several common infectious diseases of the 1950s, and results show geographical equilibrium distributions for several epidemics. These are not predicted by the most commonly used epidemiological models but are consistent with observed geographical disparities in disease prevalence that continued over a number of years in spite of recurrent epidemic cycles and long-term trends.

## 1. Introduction

Mathematical models of epidemics provide great insight as to their development over time and whether they will burn out or reach an endemic equilibrium. Increasingly, researchers are giving more attention to the spatial dimension of an epidemic over time and the conditions for endemic equilibrium, which are the topics of this research. A steady-state incidence of disease on the time dimension does not necessarily imply a uniform spatial distribution. In fact, as evidence here will show, spatial differences in disease rates can persist for years even in the face of recurrent epidemic cycles. Epidemiologists generally attribute such spatial differences to heterogeneity in the population (Keeling and Rohani, 2008). For example, there may be underlying socioeconomic conditions that differ across regions, resulting in different disease rates; or the dynamics of the disease may be at different stages across a geographic area. Research here, however, will give a different explanation for this situation based on a spatial model of disease contagion or diffusion. This also leads to the conclusion that the prevalent mathematical models of disease spread do not adequately explain observed spatial distributions of diseases.

For discussion purposes one can distinguish two broad classes of models: those based on the SIR model and others. At the macro level, change over time and equilibrium conditions have been well studied analytically for the SIR model and its variants. The model uses three simultaneous, ordinary differential equations to explain the proportions of the population who, over time, are susceptible to infection (S), infected (I), or have recovered (R). The model assumes, however, that all susceptible people have an equal chance of being infected at any time, which precludes analysis of the spatial diffusion of a disease. The model must be expanded to study diffusion. Here the analysis concerns the continuous geographical distribution of common infectious diseases across the United States, and a partial differential equation model is best suited to the analysis. Typically the model is the reaction-diffusion equation. Originating in chemistry, these are widely used in ecology (Holmes, et al., 1994) and have been adapted to epidemics (Kendall, 1965; Noble, 1974; Mollison, 1977; Murray, Stanley and Brown, 1986; Lopez, et al., 1999; Reluga, 2004). They incorporate the features of the SIR model that explain the course of an epidemic over time and add the term *u_xx_ + u_yy_*, the sum of the second partial derivatives of change in two spatial dimensions, to model diffusion. Note that it is the movement of infected people that spreads the disease, not movement of the pathogen by itself. Analysts sometimes distinguish two types of diffusion: the familiar spread from one area to the next and so-called hierarchical diffusion, from cities to rural areas; but this analysis will not make the distinction.

As to whether an endemic equilibrium is reached, the SIR model with births and deaths predicts two possible outcomes (Keeling and Rohani, 2008): the epidemic dies out, or it reaches a stable fixed proportion of susceptible people equal to the inverse of the basic reproductive rate–the average number of people infected by a sick person when no one has immunity. The steady-state proportion of the population who are infected depends on the reproductive rate, the transmission rate of the infection, and the natural death rate. Continued oscillations can result, however, owing to seasonal forcing, such as when children go back to school in the fall (Keeling and Rohani, 2008), or when imperceptible changes in the transmission rate are amplified dramatically by dynamic resonance (Dushoff, et al., 2004).

Reaction-diffusion models add substantial complexity to the SIR model and to questions about equilibrium. In the simple case of a single source of infection the model is easily solved and predicts a travelling spatial wave moving out from the source. More complicated situations can lead to intractable solutions. However, because partial differential equations can be solved numerically using a discrete lattice model as an approximation, the behavior of a reaction-diffusion equation is often studied by a computer simulation of the corresponding lattice model. Ultimately an infection may die out or reach an endemic equilibrium, as one might expect from the SIR model, or not, as Liu and Jin (2007) show using computer simulation of a lattice model. Complex labyrinthine spatial patterns can arise when births and deaths play a role, and different values of parameters lead to stability or instability. Often these mathematical models are not explicitly geographical, that is, applied to location data for an actual country, or they are applied to relatively simple cases. But Cahill et al (2009), for example, analyze influenza in the U.S. at a very fine spatial detail, combining partial differential equation models in their discrete form with computer simulation and actual data.

Non-SIR approaches also are used to study epidemics as a continuous function of location. These usually focus on the spread of disease across an actual geographic area but are less suited to studying conditions of an equilibrium analytically. Grenfell, Bjornstad, and Kappey (2001) demonstrate spatial waves in measles epidemics in the U.K. with a wavelet analysis of weekly morbidity data from the 1950s and 1960s. Treveleyan and Smallman-Raynor (2005) make an exhaustive analysis of polio epidemics in the U.S. from 1910 to 1971 using a spatial autocorrelation measure (Moran’s *I*) and time-series analysis. Pyle (1986) and Xia et al. (2004) are among the analysts who use a gravity model, for which the diffusion between two areas is proportional to the product of their populations and inversely related to the distance between them to (perhaps to a power). Xia applies this model to measles epidemics in the U.K. comparing a computer simulation with actual data.

This analysis combines geographical data on disease rates with a partial differential equation model for which there is an analytical solution to the equilibrium question. The model is applied at the state level to a variety of infectious diseases that were prevalent in the U.S. in the early 1950s. The central questions are whether there is evidence of a spatial equilibrium in certain endemic illnesses, and whether certain epidemics at times may have briefly reached an equilibrium across the U.S. despite annual cycles, long-term trends, and geographically sporadic outbreaks. Was there an endemic spatial equilibrium underneath the epidemic cycles? Historical evidence suggests this possibility. For example, among the endemic illnesses, syphilis and TB had a long history of high rates in the Southeast and Southwest, respectively, and lower rates in the North. This relates to social conditions among nonwhite populations in those regions. Moreover, historical evidence for epidemic illnesses also shows that boundary states tended to have extreme values and that geographical disparities persisted from year to year despite annual epidemics. Regional diphtheria rates from 1948 to 1954, for instance, display consistent differences between regions, with the highest levels in the East South Central and West South Central and lowest infection rates in the Middle Atlantic states (Moore and Larsen, 1957). From the 1940s through 1950s polio, too, had distinct geographic patterns, which gradually changed over time (Serfling and Sherman, 1953). From 1947 to 1951 geographic areas west of the Mississippi had considerably higher rates than those to the east. The reasons for this are not known.

The focus is on several diseases that are transmitted mainly by personal contact and are highly contagious, so that a diffusion process is clearly involved in their spread. These are diphtheria, polio, measles, whooping cough (pertussis), tuberculosis, and syphilis. All these illnesses except polio were in a general decline by the 1950s but still common. Total reported cases in 1950 were: measles 319,000; syphilis (1952) 169,000; TB 122,000; whooping cough 121,000; polio 33,300; and diphtheria 5,800. National and state data on disease incidence was collected through a mandatory reporting system that was well developed by the 1950s, but completeness of reporting cannot be guaranteed. Whether the disease reached a clinical state also bears on reporting; with polio, in particular, 95 percent of infections did not reach clinical significance. Vaccination against diphtheria had been available by the 1950s, but it was not yet widespread enough to prevent yearly outbreaks (Moore, 1958), while the Salk vaccine did not undergo field testing until 1954. Polio, diphtheria, and measles epidemics recurred in large annual cycles, whereas whooping cough was less variable over time, and TB and syphilis were endemic but subject to local outbreaks. With TB, in fact, people can develop the illness even years after exposure.

## 2. Model

In its simplest form, the mathematical model for spatial diffusion over time of a quantity *u* in two spatial dimensions is

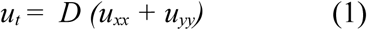

Ideally one would model the spread of diseases with Equation (1) using both time series and cross-sectional data, but this has severe practical difficulties. The extent of complete time series data is limited. Before the Second World War state reporting was incomplete, and by the late 1950s the diseases were in significant decline owing to antibiotics, vaccination, and public health measures, meaning that a partial differential equation model would have to be time varying in its coefficients and therefore very complex—too complex for the available data. So the approach here is limited and modest but offers a snapshot of the geographical distributions within a limited and relatively stable regime not to be repeated again.

As the diffusion process evolves over time, eventually a steady state is reached, when *u_t_* = 0, giving the Laplace equation (2)

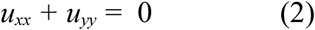

Consider that one knows the value of *u(x,y)* on a two-dimensional lattice of points at a small equidistance *h* from one to the next with lattice representing a geographical area. One can approximate Equation (2) on the lattice at point *(m,n)* by a Taylor expansion about *(x,y)*, which after dropping negligible terms leads to

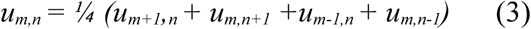

So the value of *u* at each point on the lattice is approximately the average of the values at the four adjoining points on each side. This approximation, Equation (3), can be used to solve the equation numerically through an iterative, relaxation process.

Let us express the idea that because of contagion each unit becomes more like its neighbors, with prevalence rate *u* of a disease at *(x_i_*, *y_j_)* tending toward the average rates in the four neighbors. The units might have any rates initially. One can extrapolate what will happen in this arrangement by a mental or computer simulation. At each iteration one successively replaces the rate at each point by the average rate of its four neighbors.

Given observed spatial disparities, it is also assumed that disease rates in the units on the geographic boundary of the country (or lattice) do not change, or at least change very slowly in relation to change in the interior. This agrees with historical evidence for several diseases and is reasonable theoretically because each boundary unit interacts with two neighbors that are also boundary units but with only one interior unit. Change in the interior propagates slowly to the boundary. It is assumed that no contagion occurs across the borders.

This model (3) leads to a distribution across the country or lattice that is unique and depends only on the values on the boundary. If the simulation continues until no further change occurs—the steady state—the distribution of the disease fits a mathematical function *u(x,y)* known as a harmonic or potential function (Garabedian, 1964: 458ff). It is this type of function that interests us, not the actual values. Such a function is a unique solution of the Laplace equation (2).

One can compare the diffusion of a disease by analogy with heat diffusion as seen, for example, in the daily weather maps that show contours of temperature across the country. So as a standard of comparison with the an endemic equilibrium the analysis here also includes a model for the distribution of average state temperatures across the 48 states from 1971 to 2000.^1^ The averaging reveals the climate equilibrium underneath seasonal cycles and other short-term temperature fluctuations (neglecting any mention of global warming). By analogy, the analysis here seek to uncover and explain any underlying steady-state in the geographic distribution of diseases, within the limits of the data.

A harmonic function has unique properties (Kellogg, 1953) that one can use to test for the distribution: (1) The product of a harmonic function multiplied by a constant is harmonic (scale invariance), as is the sum or difference of two such functions. (2) It is invariant— still harmonic—under translation or rotation of the axes. (3) The function over an area is completely determined by the values on the boundary; the solution is unique. (4) A harmonic function over a closed, bounded area takes on its maximum and minimum values only on the boundary of the area (if it is not a constant). (5) If a function is harmonic over an area, the value at the center of any circle within the area equals the arithmetic average value of the function around the circle. This implies that averages around concentric circles are equal. The converse is also true. If the averages around all circles equal the values at their centers, the function is harmonic. Note also that the property of scale invariance implies that the size of the units of analysis should not matter much.

Here three properties of harmonic functions are tested to validate the model: (1) that the geographical distribution of an epidemic disease is a harmonic function; (2) that average rates around concentric circles are equal; and (3) that the maximum and minimum disease rates are in border states. These hypotheses would be satisfied trivially if the distribution is constant, so from a conservative approach to hypothesis testing, this situation must be ruled out as well. And one must verify that the distribution in not random. The specific harmonic function that concerns us here is a plane surface that is a function of latitude and longitude. A broad class of nonharmonic alternatives to this can be tested statistically with quadratic equations, such as *u(x,y)* = *a x*^2^ + *b x* +*c* or *u(x,y)* = *a x*^2^ + *b x y* + *c y*^2^ + *d* when *a + b + c* ≠ 0. The analysis is limited, however, to testing these hypotheses with areal data, which has a high degree of granularity as to location. So the hypotheses must be adapted to fit this type of data.

## 3. Spatial Analysis

The research plan was to examine the disease rates at approximately the same time period to allow a simultaneous comparison of their geographical distributions and possible equilibria. For diphtheria, two data elements are included: the state average rates from 1950 to 1954 (Moore and Larsen, 1957), and the 1955 rate (Moore, 1958). State measles and whooping cough rates are for 1950 (Vital Statistics of the United States 1950 Vol.1, 1953). Polio data includes 1949, 1950 (Dauer, 1951) and, separately, the first quarter of 1950 (Public Health Reports,1950) which was the low point of the epidemic between epidemics of 1949 and 1950. TB data are an average from 1949 to 1951 (Anderson and Sauer, 1952), which helps to smooth out minor temporal and local fluctuations, and the natural logarithm of TB rates was used to further moderate the effect of some extreme values. Syphilis rate data is for fiscal year 1952 (Public Health Reports, 1954). Availability of data influenced these choices as well.

This analysis uses the geographical software GeoDa 0.9.5 developed primarily by Luc Anselin, who pioneered many of the methods used in spatial analysis. GeoDa follows the ArcView standard for geometric area data files developed by ESRI, Inc. To construct a map and analyze the corresponding data, three different files are required: a shape file (*.shp) that describes the geometry of each unit, an index file (*.shx), and a data file in dBase (*.dbf) format. GeoDa is available at no charge via the Internet from Arizona State University.^2^

### 3.1 Test for spatial autocorrelation

The first task is to check that the spatial distribution of each disease is not random. The spatial autocorrelation is the correlation between the rate in each state and the average of rates in neighboring states. So this is also a first indication of diffusion of an illness among neighboring states. For this analysis the neighbors around each state are the set of states that have a boundary in common with it, which is called rook contiguity by analogy with chess. This is a gross approximation of the lattice model discussed earlier but is sufficient to begin testing the model. In the U.S. this identification of neighbors leads to different numbers for the states.^3^ The most common number of neighbors is four, and forty states have between three and six states sharing a border.

Spatial autocorrelation for the entire country is assessed with Moran’s *I*. This is a measure of spatial autocorrelation with range [−1,1]. As with Pearson’s correlation, Moran’s *I* can be positive or negative, and a value close to zero implies no autocorrelation. It is based on the aggregate of autocorrelations in the neighborhoods of all states. When states with above average disease rates are neighbors of states that also have above average rates, the *I* value increases; the same holds when below average states border other below average states. A seen in Table 1, all diseases but whooping cough are statistically significant on this measure (at p < .05); whooping cough narrowly misses the cutoff at p = .07. Diseases showing the strongest correlation between a state’s infection rate and the average of its neighbors are, in decreasing order of strength: average diphtheria, polio in the first quarter of 1950 and in 1949, diphtheria in 1955, syphilis, TB, measles in 1950, and polio in 1950. Significance levels are determined by a permutation test.

**Table 1.**
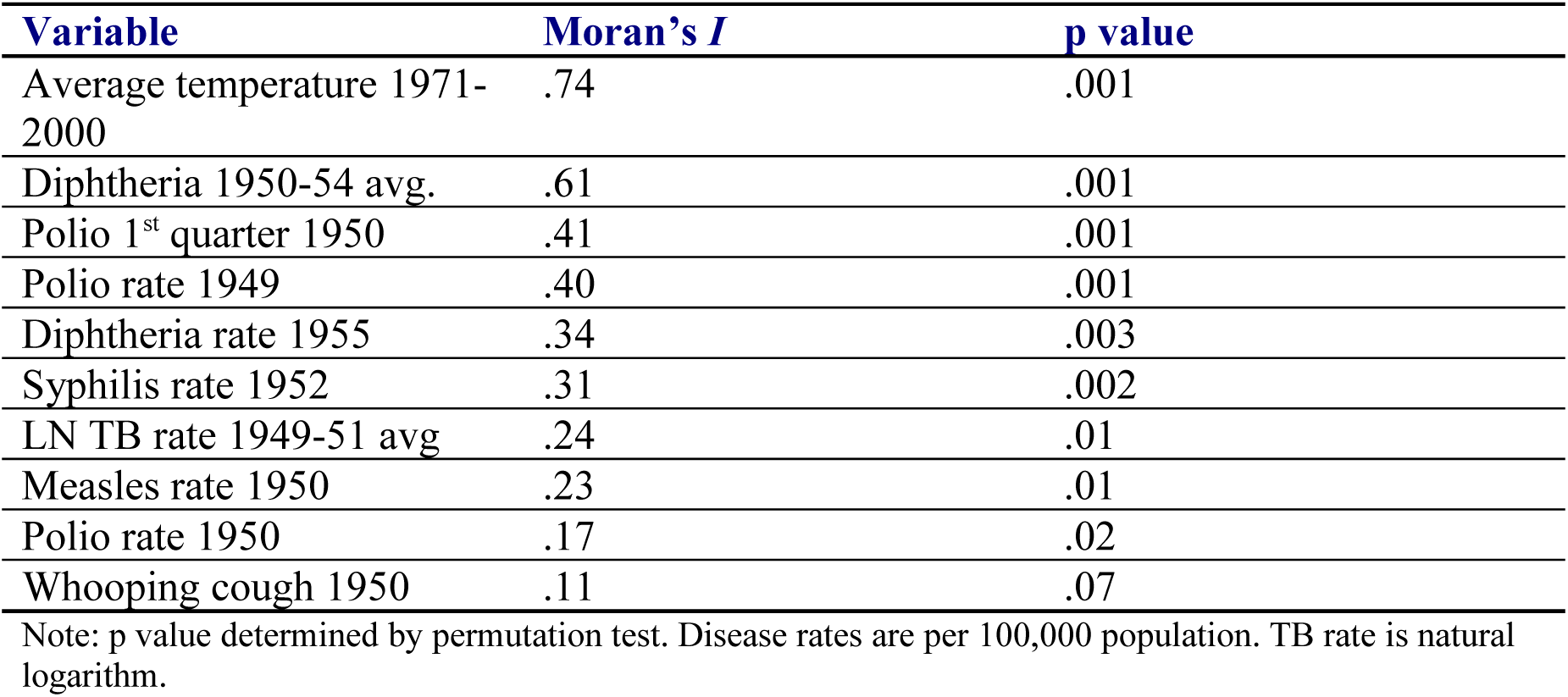
Moran’s test for spatial autocorrelation.

| **Variable** | **Moran's *I*** | **p value** |
| --- | --- | --- |
| Average temperature 1971-2000 | .74 | .001 |
| Diphtheria 1950-54 avg. | .61 | .001 |
| Polio 1<sup>st</sup> quarter 1950 | .41 | .001 |
| Polio rate 1949 | .40 | .001 |
| Diphtheria rate 1955 | .34 | .003 |
| Syphilis rate 1952 | .31 | .002 |
| LN TB rate 1949-51 avg | .24 | .01 |
| Measles rate 1950 | .23 | .01 |
| Polio rate 1950 | .17 | .02 |
| Whooping cough 1950 | .11 | .07 |

Note: p value determined by permutation test. Disease rates are per 100,000 population. TB rate is natural logarithm.

### 3.2 Harmonic function test

If the variable distributions are solutions to the Laplace equation, they should be harmonic functions of their location. In this case, the models represent the equilibrium state resulting from a nationwide spread of the epidemic. Inspection of maps of the state distributions suggests that one can try to model the distribution as a function of latitude and, possibly, longitude. The map shapefile contains information on the longitude and latitude of the vertices of the polygon used to map each state. For each state GeoDa can compute a centroid, which is the latitude-longitude location of the geometric center of gravity of the state. This location is used in the analysis. Table 2 shows the results of linear regression of each disease distribution and average temperature against latitude or longitude (and those variables squared if statistically significant) at the state centroid.

**Table 2.** National model for harmonic function hypothesis test: *Y* = constant + *B_1_* * Latitude + *B_2_* * Latitude^2^ + *B_3_* Longitude + *B_4_* Longitude^2^

| Variable | Constant<br>(error) | $B_1$<br>(error) | $B_2$<br>(error) | $B_3$<br>(error) | $B_4$<br>(error) | R<br>square | p | LM<br>spatial<br>lag p | LM<br>spatial<br>error p |
| --- | --- | --- | --- | --- | --- | --- | --- | --- | --- |
| Average<br>Temp. | 111 (3.2) | - 1.50<br>(0.08) | ns | ns | ns | .88 | <.0001 | .18 | .006 |
| Syphilis<br>1952 | 555 (60) | -11.5<br>(1.5) | ns | ns | ns | .56 | <.0001 | .09 | .03 |
| Diphtheria<br>1950-54<br>average | 48.7<br>(15.1) | - 2.12<br>(0.78) | 0.0237<br>(0.010) | ns | ns | .44 | <.0001 | .004 | .002 |
| Number<br>Diphtheria<br>1950-54<br>average | 51,400<br>(8540) | -1120<br>(215) | ns | ns | ns | .37 | <.0001 | .72 | .92 |
| Polio 1<sup>st</sup><br>quarter<br>1950 | -3.5<br>(0.94) | ns | ns | -0.049<br>(0.010) | ns | .34 | <.0001 | .90 | .74 |
| Polio 1949 | -57 (19) | 2.25<br>(0.50) | ns | ns | ns | .30 | <.0001 | .04 | .01 |
| Whooping<br>Cough<br>1950 | 1620<br>(440) | ns | ns | 32.2<br>(9.4) | 0.17<br>(0.05) | .22 | .003 | .57 | .48 |
| LN TB<br>1949-51<br>avg. | 6.18<br>(0.56) | -0.048<br>(0.014) | ns | ns | ns | .20 | .001 | .67 | .73 |
| Diphtheria<br>1955 | 8.5 (2.9) | -0.18<br>(0.07) | ns | ns | ns | .16 | .005 | .02 | .03 |
| Number<br>Diphtheria<br>1955 | 152,000<br>(46000) | -7010<br>(2395) | 81.4<br>(30.7) | ns | ns | .37 | <.0001 | .96 | .52 |
| Measles<br>1950 | - 170<br>(218) | 9.6<br>(5.5) | ns | ns | ns | .04 | .09 | .08 | .09 |
| Polio 1950 | 17.2<br>(11.4) | 0.077<br>(0.288) | ns | ns | ns | .002 | .79 | .10 | .10 |

Notes: Latitude and longitude are at the state's centroid. The quadratic surface for average diphtheria has a minimum at 45 degrees latitude; for number diphtheria in 1955, at 43 degrees latitude; for whooping cough, there is a minimum at -97 degrees longitude. Spatial lag model (with std. errors) including lagged dependent variable for average diphtheria is $Y = 6.33 (2.21) - 0.13 (.05) \text{latitude} + 0.54 (.14) \text{spatial lag } Y$ , $R \text{ square} = .55$ . For diphtheria in 1955 the spatial lag model is $Y = 5.82 (2.53) - 0.12 (0.06) + 0.36 (.17)$ ; $R \text{ square} = 0.26$ .

Rates for diphtheria in 1955, ln TB, polio in 1949, and syphilis are linear functions of their latitude at the centroid (Fig.1–2). These are harmonic functions of latitude, namely, plane surfaces. The same applies to polio in the first quarter of 1950, which is a linear function of longitude (Fig. 3). But the average diphtheria rate form 1950 to 1954 is a quadratic surface as a function of latitude (Fig. 4), and whooping cough is a quadratic function of longitude (Fig. 5), which are not harmonic functions. The quadratic surface for diphtheria rate is close to linear over most of its range, however, having a minimum at 45 degrees latitude (the latitude of Minneapolis). The change in polio models from 1949, which had a great epidemic, to the first quarter of 1950 when the epidemic had receded almost completely, to the full 1950 epidemic is striking and not well explained. The strongly geographically oriented distribution of 1949 is gone by the full 1950 epidemic. The models also were tested with the inclusion of state population and population density in 1950; in no case were these variables statistically significant—an important point to be discussed later.

**Figure 1.**
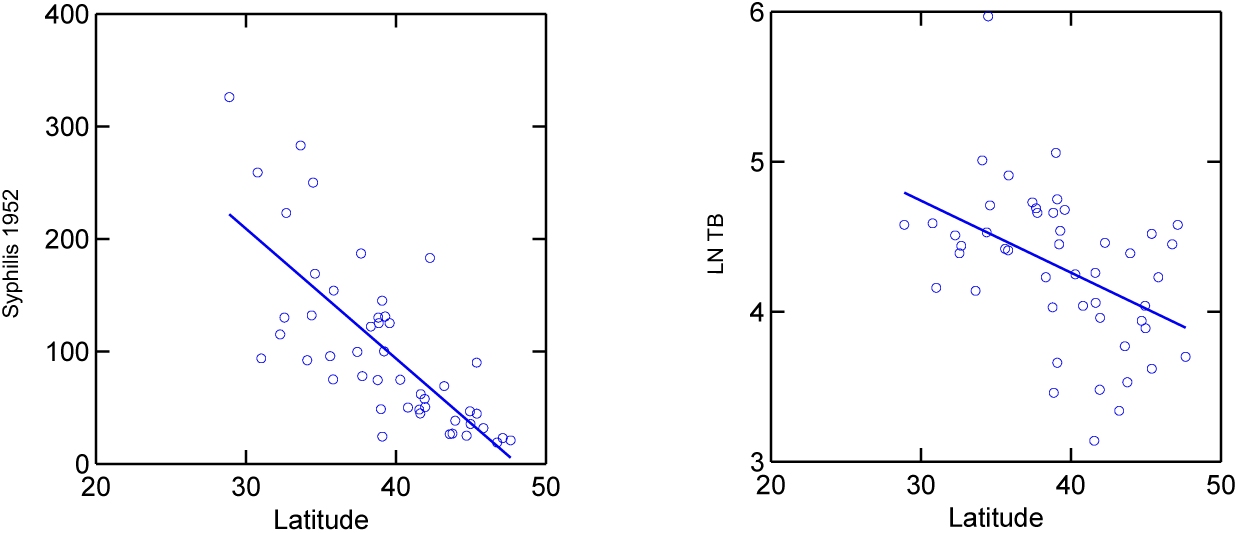
Syphilis and ln TB rates in relation to latitude with linear fit.

**Figure 2.**
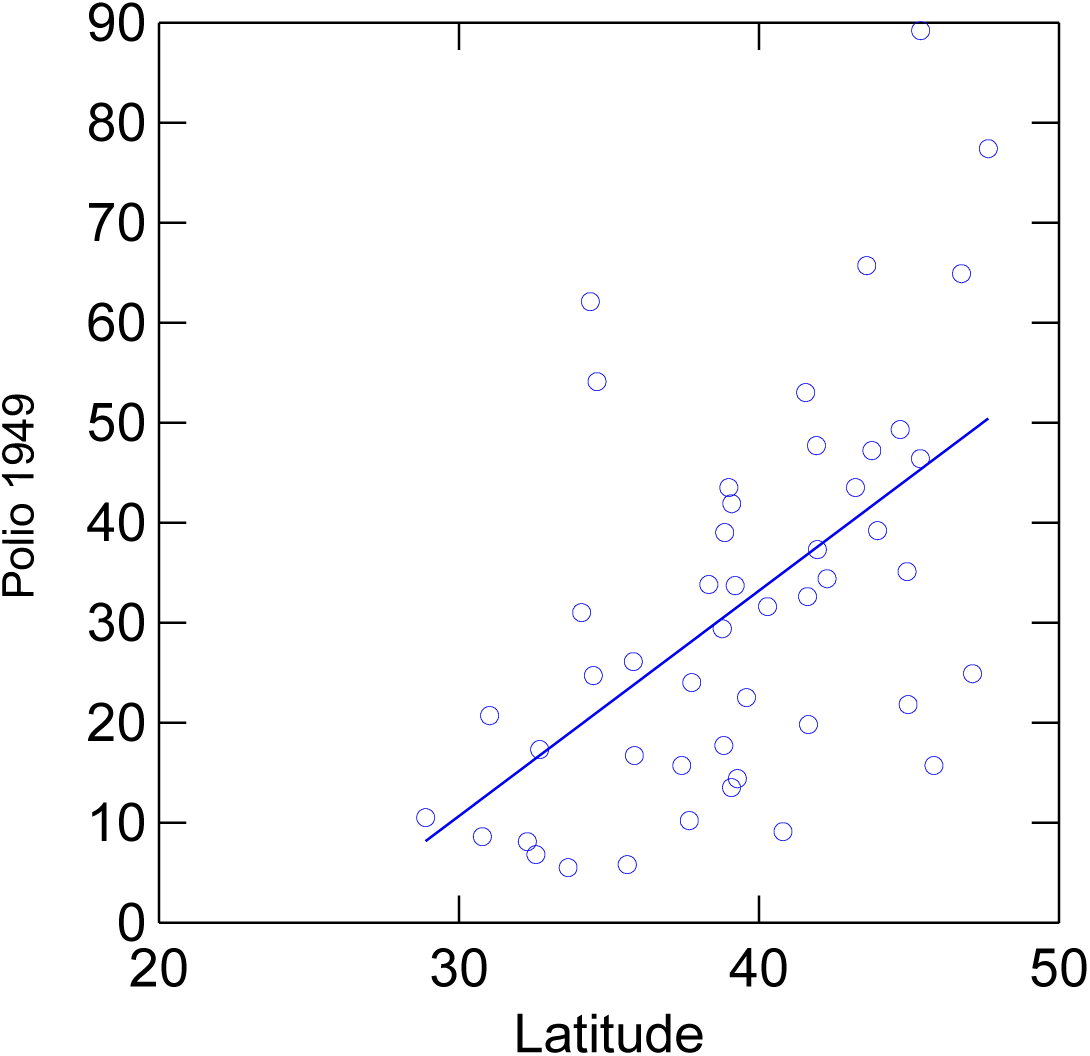
Polio rate in 1949 in relation to latitude with linear fit.

**Figure 3.**
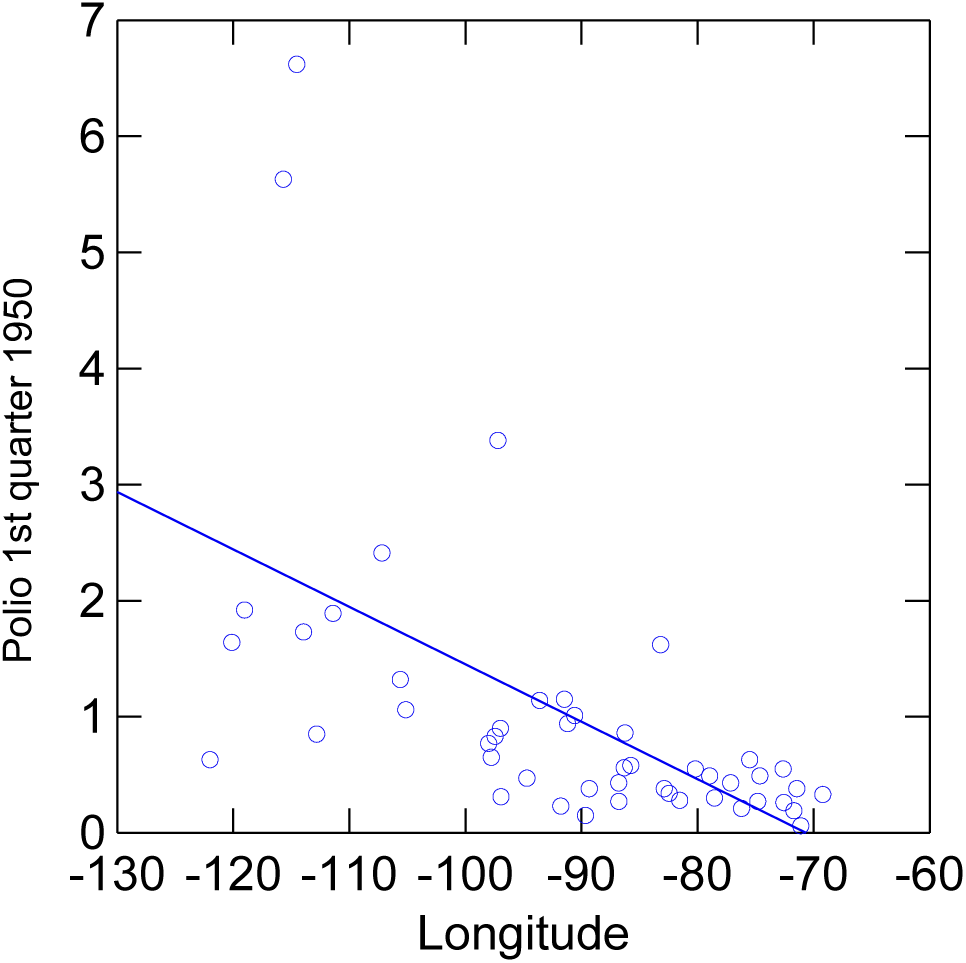
Polio rate in the first quarter of 1950 in relation to longitude with linear fit.

**Figure 4.**
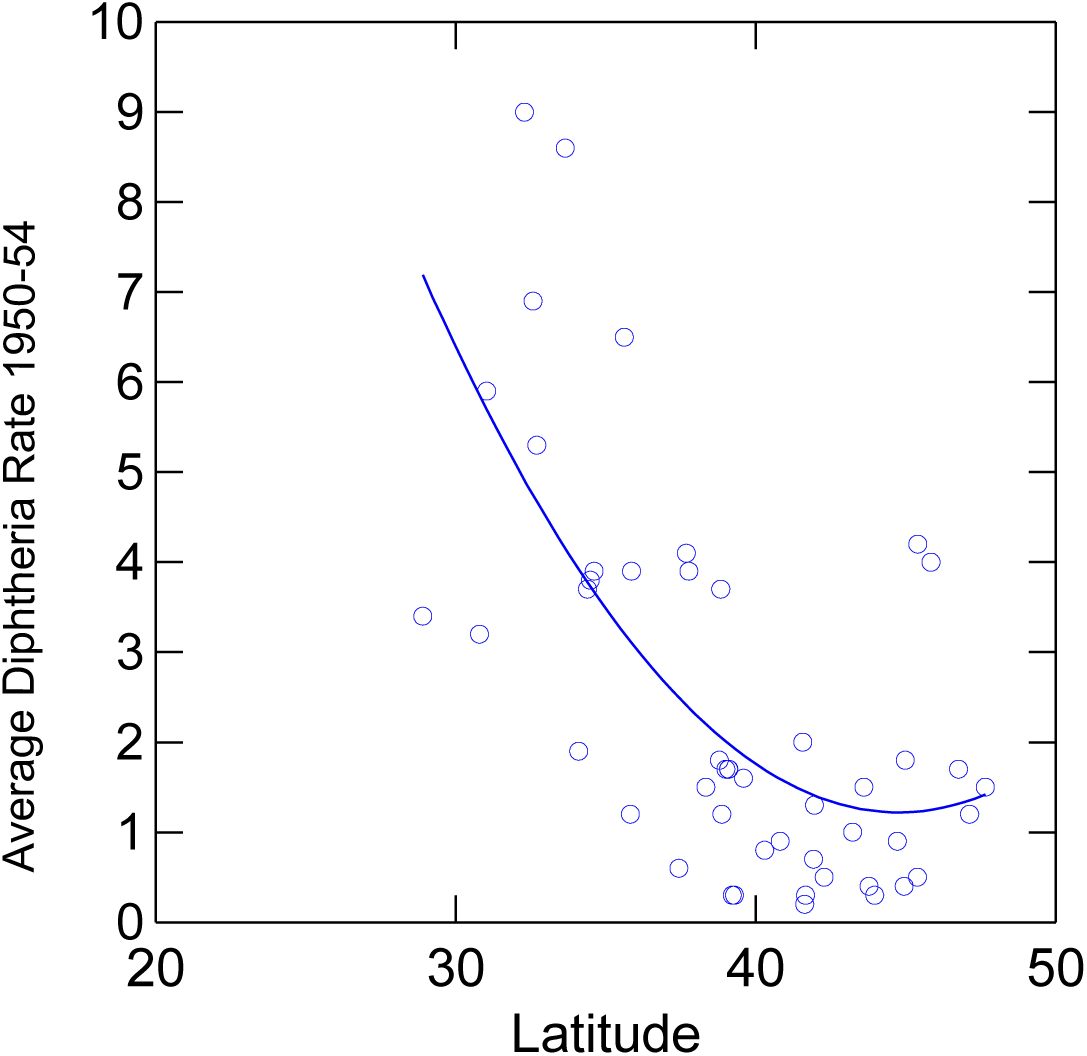
Diphtheria rate in relation to latitude; 1950–54 average with a quadratic fit.

**Figure 5.**
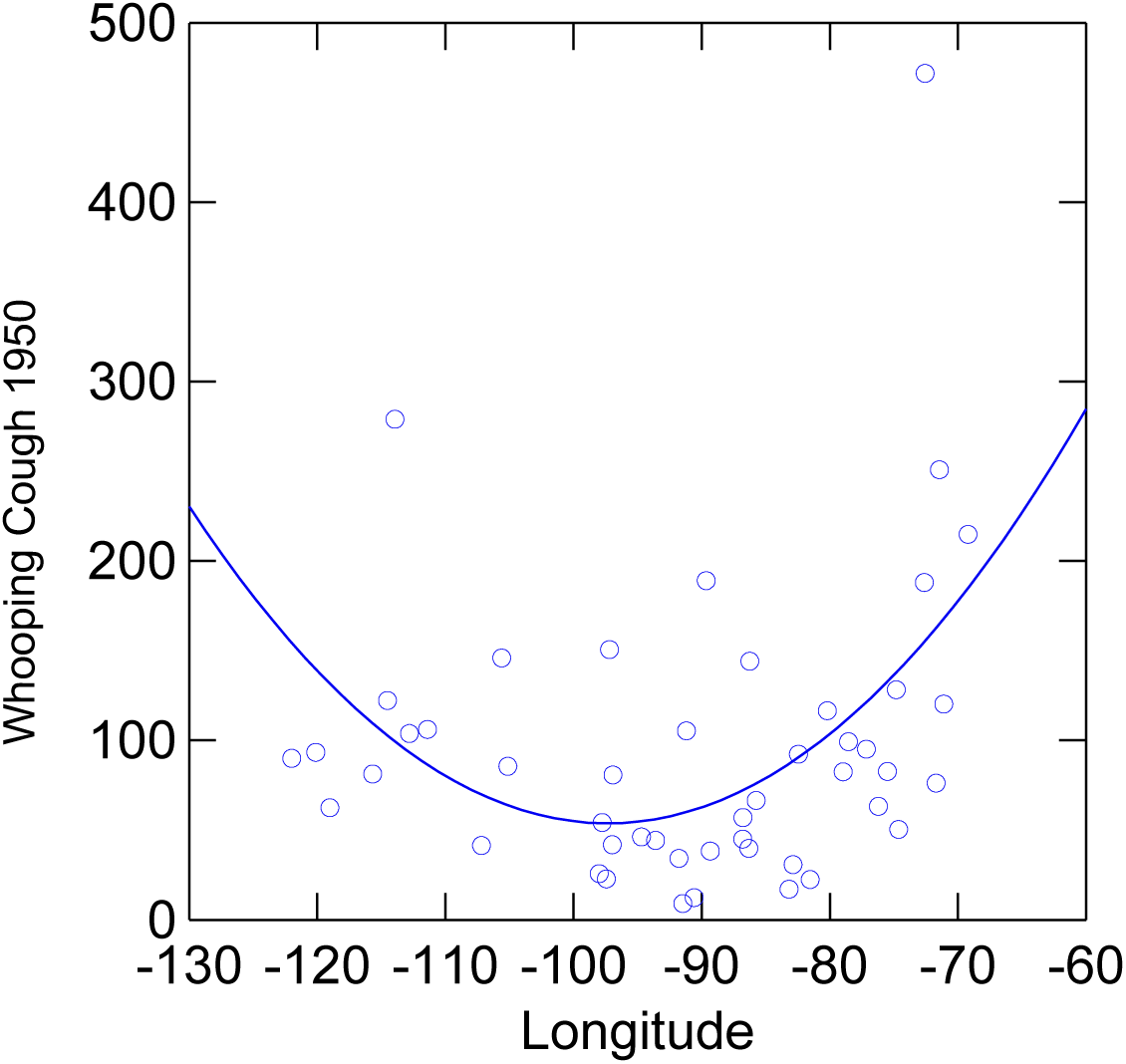
Whooping cough rate in 1950 in relation to longitude with quadratic fit.

The analysis checked whether the latitude-longitude models capture all the spatial lag. As seen in Table 2, Lagrange Multiplier (LM) tests indicate models where there is remaining spatial lag and whether there is error correlation between units. When spatial lag is significant, the OLS coefficient estimate is biased and usually too large. The LM tests may also indicate a missing variable, inefficient estimates, or other regression problems. The spatial lag of the dependent variable remains an issue with diphtheria; for other diseases the effects are marginal or nonexistent. One can include the lagged dependent variable in the regression (note, Table 2). Such a model would suggest that the diphtheria rate in a state depends on both its latitude and the rates in adjacent states, and that the distribution had not reached an equilibrium, although the spatial lag term weakens by 1955. Further analysis, however, indicates an alternative approach as a model for diphtheria at equilibrium.

Recall that in 1950 there were only 5,800 diphtheria cases reported nationally, and several states had no cases. This number is very small in comparison to the other epidemics studied. In such a situation it may be that the actual number of cases is more important than the rate in determining the spread of a disease. (This is often the case for animal diseases in the wild where populations are relatively small.) To test this idea, the analysis of diphtheria was redone with the average number of cases from 1950 to 1954 and the number in 1955. Results (Table 2) show that this procedure completely eliminates any remaining effect of spatial lag or error; only latitude is statistically significant. But the model for 1955 is a quadratic function of latitude while the average number is a linear function (Fig. 6) (just the reverse of the OLS model for rates.) Generally, however, the shapes of the distributions on both models are similar over most of the range of latitude. The second approach, however, has the advantages of eliminating any problems related to unexplained spatial effects, giving an accurate estimate of the latitude coefficients in the simpler OLS model, and confirming the prediction.

**Figure 6.**
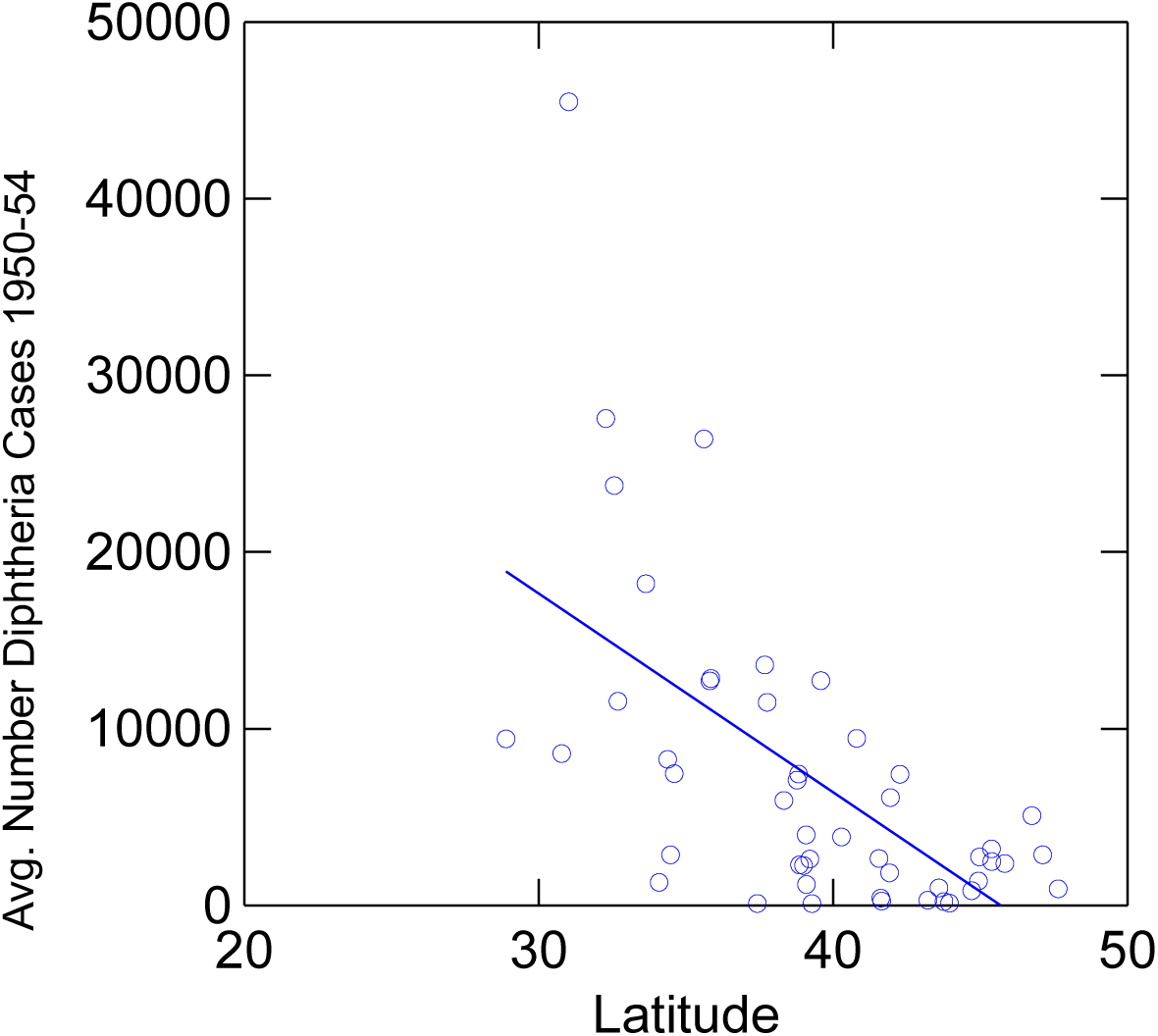
Average number of diphtheria cases 1950–54 in relation to latitude with linear fit.

Because a harmonic distribution is completely determined by its boundary values, the analysis was redone for those 30 states only.^4^ Results are in Table 3. A comparison of models in Table 2 and 3 shows that for diphtheria (without spatial lag), syphilis, measles, and polio in 1949 the coefficients for latitude are almost the same, excepting quadratic models. The correspondence is somewhat weaker for TB. Whooping cough has the same nonharmonic model in each. Where the two models give about the same results, the boundary values, therefore, are about as accurate in prediction of the values of interior states as is the model for all states. This is further evidence that the spatial distribution is close to an equilibrium and in accord with the assumption of fixed boundary values.

**Table 3.** Boundary states model for harmonic function hypothesis test (N = 30): *Y* = constant + *B_1_* * Latitude + *B_2_* * Latitude^2^ + *B_3_* Longitude + *B_4_* Longitude^2^

| **Variable** | **Constant**<br>**(error)** | **$B_1$**<br>**(error)** | **$B_2$**<br>**(error)** | **$B_3$**<br>**(error)** | **$B_4$**<br>**(error)** | **R**<br>**square** | **p** |
| --- | --- | --- | --- | --- | --- | --- | --- |
| Avg. temp. | 111 (3.3) | - 1.48<br>(0.08) |  |  |  | .92 | <.0001 |
| Syphilis<br>1952 | 560 (76) | -11.4<br>(1.9) |  |  |  | .56 | <.0001 |
| Polio 1949 | -56.1<br>(16.8) | 2.10<br>(0.42) |  |  |  | .47 | <.0001 |
| Diphtheria<br>1950-54<br>average | 14.3 (2.6) | -0.30<br>(0.07) |  |  |  | .42 | .0001 |
| Number<br>diphtheria<br>1950-54<br>average | 54,400<br>(1,100) | -1170<br>(278) |  |  |  | .39 | .0002 |
| Diphtheria<br>1955 | 10.7 (2.7) | -0.23<br>(0.07) |  |  |  | .30 | .002 |
| Number<br>diphtheria<br>1955 | 35,800<br>(8,400) | -786<br>(211) |  |  |  | .33 | .001 |
| Polio 1<sup>st</sup><br>quarter<br>1950 | 16.5 (6.2) | -0.77<br>(0.33) | 0.0092<br>(0.0042) |  |  | .26 | .006 |
| LN TB<br>1949-51<br>avg. | 5.56 (0.50) | -0.029<br>(0.013) |  |  |  | .16 | .02 |
| Whooping<br>Cough<br>1950 | 1711 (647) |  |  | 33.7<br>(13.9) | 0.17<br>(0.07) | .13 | .05 |
| Measles<br>1950 | - 162<br>(205) | 9.0 (5.2) |  |  |  | .10 | .09 |
| Polio 1950 | not<br>significant | ns |  |  |  | 0 | ns |

Notes: Latitude and longitude are at state's centroid. Quadratic surface for polio 1<sup>st</sup> quarter has a maximum at 42 degrees latitude; whooping cough surface minimum is at - 99 degrees longitude.

### 3.3 Mean-value test for concentric circles

The next analysis checks the hypothesis that mean values around a circle equal the value at the center. As an approximation, one can compare the average value of the boundary states with that of interior states. A t-test shows that for syphilis, rate and number of average diphtheria 1950–54, rate and number of diphtheria in 1955, polio in the first quarter of 1950, polio in all of 1950, and measles in 1950 there is no statistically significant difference in mean rates of infection between boundary and interior states. Differences are statistically significant for polio in 1949, whooping cough, and ln TB. One must be cautious about interpreting the results, however, when the distribution is close to uniform across the country, subject to random variation at the state level. In such a case one would be likely to find no difference in means. This might be the situation with polio and measles in 1950, which do not have a statistically significant relationship with latitude or longitude. In sum, diphtheria, syphilis, and polio in early 1950 best meet the test of this hypothesis.

This proposition also was tested with a nonparametric Wilcoxon-Mann-Whitney test, which compared the ranked values for boundary and interior states. A statistically significant difference indicates that one of the two distributions is shifted in relation to the other. This test gives exactly the same results for each illness as the t-tests.

### 3.4 Maximum and minimum test

The maximum and minimum for a harmonic function should be on the border. Table 4 shows the states where these occur, omitting the diseases already shown not to be harmonic functions. Excepting also diphtheria rates, out of 6 cases with 12 predictions there are three missed predictions: Idaho (twice for polio) and Nebraska (TB). (Recall that Idaho was classified as an interior state because of its relatively short international boundary.) The probability of a maximum or minimum being in a boundary state is 30/48 = 0.625, so by the binomial theorem the chance of obtaining exactly 9 correct predictions of 12 is p = .17. One might say the hypothesis is supported by a preponderance of evidence but not beyond a reasonable doubt. Not all cases are statistically independent of one another, however.

**Table 4.** Locations of state maxima and minima.

| **Variable** | **Maximum** | **Minimum** |
| --- | --- | --- |
| Average temperature | Florida | North Dakota |
| Average diphtheria rate<br>1950-54 | Alabama | Connecticut |
| Average number diphtheria<br>cases 1950-54 | Texas | Delaware |
| Diphtheria rate 1955 | Alabama | ME,VT,RI,CT,ID,WY (tie) |
| Number diphtheria cases<br>1955 | Alabama | ME,VT,RI,CT,ID,WY |
| Syphilis 1952 | Florida | Minnesota |
| TB | Arizona | Nebraska |
| Polio 1949 | Idaho | South Carolina |
| Polio 1<sup>st</sup> quarter 1950 | Idaho | Massachusetts |

## 4. Conclusion

The analysis shows that several common infectious diseases of the 1950s had approximately an equilibrium in their geographical distribution across the states of the U.S. This is true for polio, TB, diphtheria (depending on the model), and syphilis. Their spatial distributions have the characteristics of a harmonic function, which is a solution to the Laplace equation, and therefore represents the end state of a diffusion process—and usually the geographical gradient was in a north-south direction, a function of latitude. This result is largely confirmed by additional tests: the regression models for boundary states, the equality of means between interior and boundary states, and the locations of maxima and minima. No analysis was made of the stability of the equilibria, but the geographic equilibria for polio are clearly unstable.

The harmonic function models allow us to explain spatial distributions across the U.S. without further consideration of heterogeneity in state populations. The only role for heterogeneity in these models is that high levels of syphilis and TB were fostered by social conditions in the Southeast and Southwest, respectively. Any remaining effects of state-level heterogeneity are in the unexplained variation of the regression model. These would include other features of geography, topography, social linkages, and so forth that might additionally affect the spread of illnesses.

Despite similar modes of infection and similarly susceptible populations, the childhood diseases do not reveal many commonalities in their geographical distributions. For measles, the main indicator of a geographical contagion effect is Moran’s *I*, but the relation between a state’s infection level and the average in neighboring states is relatively weak. This degree of spatial autocorrelation for measles was not great enough to span the country uniformly, which might have led to a harmonic function distribution. Unlike polio and diphtheria, whooping cough did not have a harmonic function distribution, although it had a distinct spatial distribution, a quadratic surface. Clearly there was something quite different about the spread of whooping cough compared to the other childhood diseases. Whooping cough also differs in not having large annual cycles of infection, but overall the results suggest that the large annual cycles one sees for polio, measles, and diphtheria are neither necessary nor sufficient for a spatial equilibrium. State population heterogeneity seems unlikely to explain spatial differences between the childhood diseases; it is hard to imagine what types of heterogeneities could have such disparate effects on different diseases. These inconsistencies in the spatial distributions of childhood epidemics are a significant puzzle that calls for further investigation.

The analysis also raises questions about models that rely on population size or density. These variables did not contribute to the models here. In the SIR-type model population size should not matter, but the lack of significance of state population size at equilibrium especially goes against a basic premise of gravity models that the rate of disease spread between two areas is proportional to the product of their populations. Furthermore, gravity models have difficulty predicting boundary values (Bharti et al., 2008), which is not an issue in this research. As to population density, prior research (Holmes, et al. 1994) relates the velocity of the spread of infection to density. If this were true for the epidemics of 1950, however, one would expect to find it statistically significant.

These findings also go beyond what SIR-type models predict and cannot be deduced from reaction-diffusion models. Consider, for example, a typical reaction-diffusion equation for spatiotemporal change. Colizza et al. (2007) describe the spread of the Black Death in Europe with equation (4), where *I* and *S* are functions of location and time (*x,y,t*)

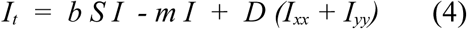

At equilibrium, *I_t_* = 0 and the equation has the general form of Poisson’s equation (5)

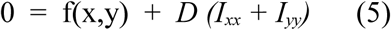

If the boundary values are held constant, as when solving the Laplace equation, the solution is unique and analogous to the surface of a drum when the skin is distorted because of a force applied to it, or it is like the distribution of temperature on the surface of a frying pan over a flame. The distribution has an equilibrium but is not a harmonic function, and with the possible exception of whooping cough, it cannot reproduce the findings seen here.

## Footnotes

1 http://www.esrl.noaa.gov/psd/data/usclimate/tmp.state.19712000.climo

2 http://geodacent.er.asu.edu

3 Because the boundary values completely determine the solution to the Laplace equation, it does not matter what the exact geometric arrangement of states is or how many share borders. This arrangement can affect the rate of convergence toward the steady-state solution, however.

4 Boundary states are: WA, OR, CA, AZ, NM, TX, LA, MS, AL, FL, GA, SC, NC, VA, MD, DE, NJ, NY, CT, RI, MA, VT, ME, OH, MI, WI, MN, IL, ND, MT. Interior states: ID, NV, UT, CO, WY, SD, OK, AR, IA, IN, KY, WV, TN, NH, PA, NE, KS, MO. States with very short national boundaries--Idaho, Indiana and Pennsylvania--are classified as interior states.

